# Blood transcriptional module repertoire analysis and visualization using R

**DOI:** 10.1101/2020.07.16.205963

**Authors:** Darawan Rinchai, Jessica Roelands, Wouter Hendrickx, Matthew C. Altman, Davide Bedognetti, Damien Chaussabel

**Affiliations:** Research branch, Sidra Medicine, Doha, QATAR; Division of Allergy and Infectious Diseases, University of Washington, Seattle, Washington, USA; Systems Immunology, Benaroya Research Institute, Seattle, Washington, USA

## Abstract

Transcriptional modules have been widely used for the analysis, visualization and interpretation of transcriptome data. We have previously described the construction and characterization of generic and reusable blood transcriptional module repertoires. The third and latest version that we have recently made available comprises 382 functionally annotated gene sets (modules) and encompasses 14,168 transcripts. We developed R scripts for performing module repertoire analyses and custom fingerprint visualization. These are made available here along with detailed descriptions. An illustrative public transcriptome dataset and corresponding intermediate output files are also included as supplementary material. Briefly, the steps involved in module repertoire analysis and visualization include: First, the annotation of the gene expression data matrix with module membership information. Second, running of statistical tests to determine for each module the proportion of its constitutive genes which are differentially expressed. Third, the results are expressed “at the module level” as percent of genes increased or decreased and plotted in a fingerprint grid format. A parallel workflow has been developed for computing module repertoire changes for individual samples rather than groups of samples. Such results are plotted in a heatmap format. The use case that is presented illustrates the steps involved in the generation of blood transcriptome repertoire fingerprints of septic patients at both group and individual levels.

## Introduction

Blood transcriptome profiling consists in measuring abundance of circulating leukocyte RNA on a global scale. This approach has been used extensively to identify changes associated with pathologies such as infection, autoimmunity, neurodegeneration, cardiovascular diseases, cancers (1–5). Another successful application has been the monitoring response to vaccines and therapeutic agents (6,7).

Reducing gene profiles to “signatures”, which are constituted of sets of genes is a strategy that is commonly employed for transcriptome data analysis and interpretation. Genes are for instance commonly grouped into sets based on similarity in expression patterns via clustering. Construction and mining of co-expression networks provides another means to identify sets of genes with similar expression profiles (8). It relies on construction and mining of co-expression networks. Nodes in such networks are constituted by genes and edges (i.e. connections) indicate gene-gene co-expression. Densely interconnected sub-networks can be extracted and constitute co-expressed gene sets also often referred to as modules.

In one such approach devised by our group the co-expression network employed for constitution of modules is based on gene-gene co-clustering. This co-clustering network is weighted to factor in co-expression of blood transcripts across multiple disease or physiological states (i.e. the maximum weigh will be attributed if, for a given gene pair, co-clustering is observed in all the states and the minimum weight will be attributed if co-clustering is observed in only one of the states).

Our first blood transcriptional module repertoire published in 2008 was based on Peripheral Blood Mononuclear Cells profiled on Affymetrix Genechips and using 8 dataset/states as input (9). It has been employed for analysis and interpretation of blood transcriptome data generated for a wide range of pathologies (9–14). A second blood module repertoire was published in 2013 (based on whole blood profiled on Illumina Hu6 Beadchips and using 9 datasets/ 7 immune sates as input (15)). The third and latest iteration is based on whole blood profiled on Illumina HT-12 chips and was developed on the basis of co-clustering observed in 16 different states, encompassing autoimmune and infectious diseases, primary immune deficiency, cancer and pregnancy (representing overall 985 unique transcriptome profiles, as described in (16)). The resulting set of 382 modules covers 14,168 transcripts. Two dimension reduction levels are built into the repertoire. The least reduced level has 382 variables, which are the individual modules (gene sets). The most reduced level has 38 variables, which are module aggregates, constituted by sets of modules. The 38 module sets encompass the 382-module repertoire. Modules have been functionally annotated through pathway, ontology as well as literature term enrichment. Module composition and high-level annotations for this third generation repertoire are available in a spreadsheet format (**supplementary File 1**). Detailed functional annotations can be accessed via interactive presentations, each corresponding to one aggregate (**Table 1**).

**Table 1:**
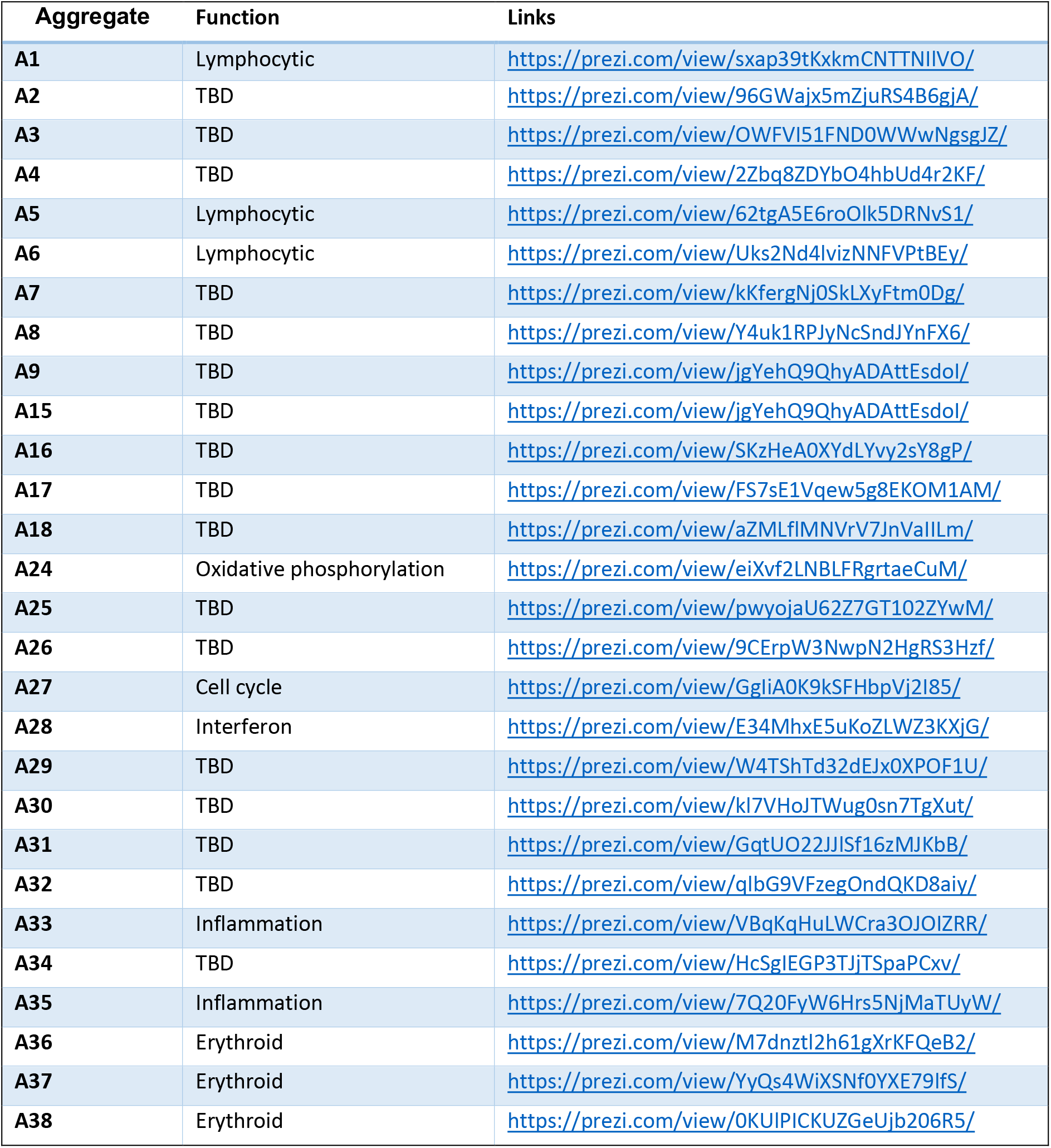
Links to module aggregates annotation pages.

Module repertoire analyses consists in determining for each module the percentage of its constitutive genes which are significantly increased or decreased. As will be described in more details below, the results of modular repertoire analyses can be represented in a “fingerprint” format, with red and blue spots indicating increase or decrease in module “activity”. These spots are represented either on a grid, with each position being assigned to a given module. They can also be represented in a heatmap format where samples are arranged in columns and modules in rows.

The script that is shared with this publication was developed to carry out these analytic steps, from annotating the gene expression data matrix with module membership information, determination of the percentage of differentially expressed transcripts among each module’s constitutive gene, to plotting those values as a fingerprint grid or heatmap.

## Results

### Preparatory steps

Here we aim to make this resource accessible to investigators who may not have an extensive bioinformatics background. In order to do so and to be very practical we chose to present the scripts that have been developed through the step-wise implementation of a use case. This section describes the initial setup phase.

First, if it is not already the case, R needs to be installed on the machine that will be used to run the analyses. If it is already installed it may also be necessary to update it to the latest version. Next, the R packages necessary to run the scripts provided as part of this publication will need to be loaded. A list of the R packages is provided in **Table 2**. They are available for download on GitHub: https://github.com/Drinchai/DC_Gen3_Module_analysis. One of the R script provided will install all the necessary packages: “*0.1Install_R_packages*”. Finally, a script is also provided to download directly from GEO gene expression matrices using GSE IDs as input: “*1.1Load_GEOdataset.R*”. It is also possible to retrieve GEO data “manually” by downloading expression files from each GEO dataset and getting the sample information from the series matrix file. This is necessary for instance when datasets comprise two different array platforms since this will prevent GEOquery from running correctly. It is the case of GSE13015, which will be used as an example in this article (17). In this step, the matrix needs is cleaned up with only the necessary information being retained. Illustrative examples are provided in **Supplementary table 1** and **Supplementary table 3** for expression matrix and sample information, respectively. This can be applied for the analysis of custom datasets.

**Table 2:** Key resources.

| Resource | Source | Ref | Identifier |
| --- | --- | --- | --- |
| GEOquery | Davis S, Meltzer P (2007) |  | https://bioconductor.org/packages/release/bioc/html/GEOquery.html |
| matrixStats v0.55.0 | Bengtsson H et. Al. (2019) |  | https://cran.rstudio.com/web/packages/matrixStats/index.html |
| gtools v3.8.1 | Warnes GR (2018) |  | https://cran.r-project.org/web/packages/gtools/index.html |
| stringr v 1.4.0 | Wickham H (2019) |  | https://cran.r-project.org/web/packages/stringr/index.html |
| ggplot2 v3.2.1 | Wickham H (2019) |  | https://cran.r-project.org/web/packages/ggplot2/index.html |
| ComplexHeatmap | Gu et al., 2016 |  | http://www.bioconductor.org/packages/devel/bioc/html/ComplexHeatmap.htm |
| preprocessCore | Bolstad B., (2020) |  | https://bioconductor.org/packages/release/bioc/html/preprocessCore.html |

For the illustrative use case presented herein the input data can be accessed as follows:

- GSE13015 expression matrix; https://www.ncbi.nlm.nih.gov/geo/download/?acc=GSE13015&format=file
- GSE13015 sample information; ftp://ftp.ncbi.nlm.nih.gov/geo/series/GSE13nnn/GSE13015/matrix/

Modular repertoire analyses consist in 4 major steps, which are depicted in **Figure 1**: 1) Annotation of the expression matrix, 2) Differential expression, 3) Calculation of the percentage of response, and 4) the visualization of module-level results (**Figure 2**: Module repertoire grid and fingerprint representation; Figure 3: heatmap). Each of these analysis steps are described in details in the subsequent sections.

**Figure 1.**
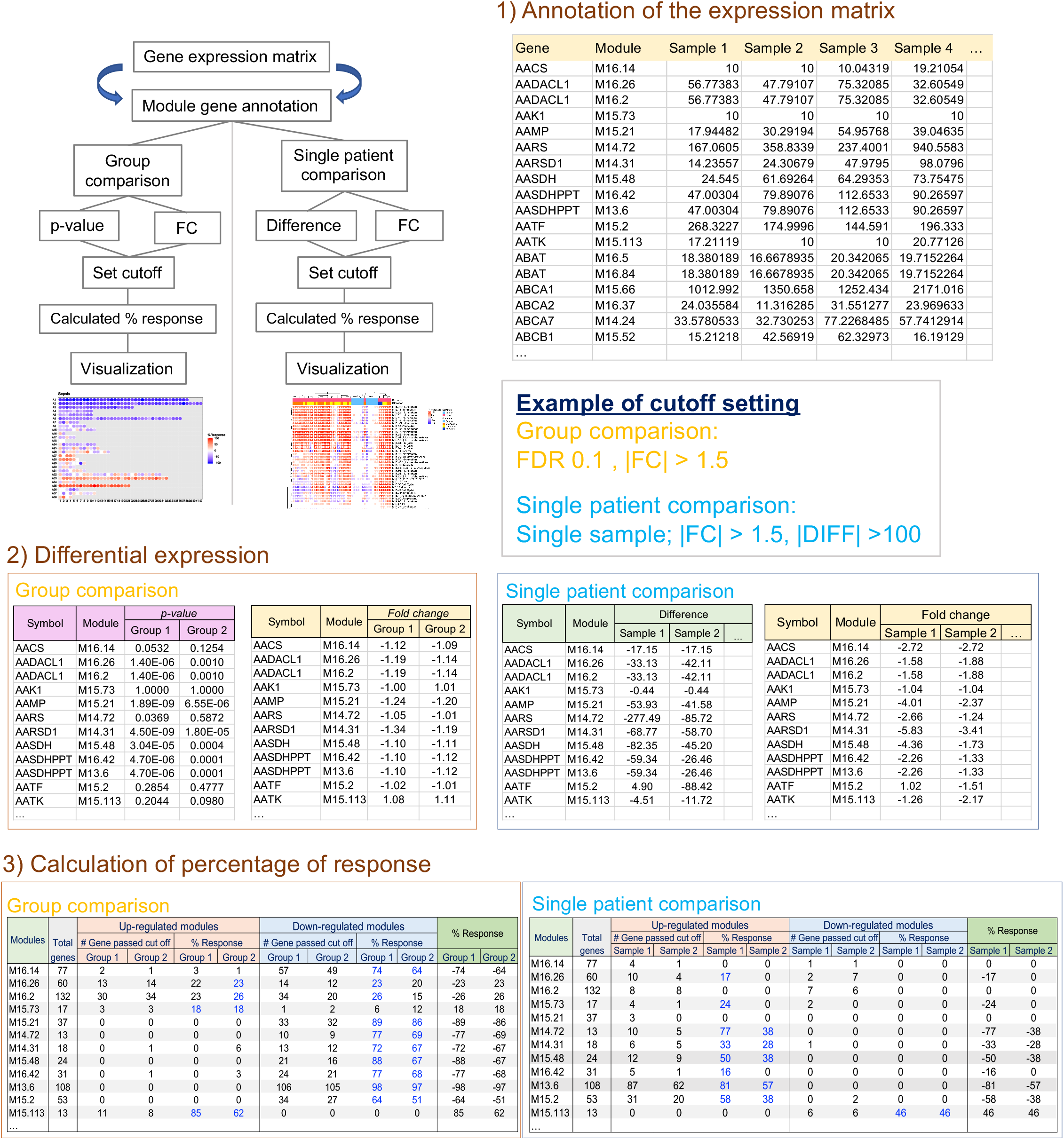
Schematic of the module repertoire analysis workflow. Briefly, steps for modular repertoire analysis include: (1) Annotation of the expression matrix: the first and second column are added to the original matrix to provide module membership information for each individual gene (Sample 1 to sample…(n)). (2) Differential expression analysis, based for instance on p-value and fold-change for comparisons at the “group level” (left panel: cases vs controls) or based on fold-change (FC) and difference at the “individual level” (right panel: individual sample vs control group). (3) Calculation of percentage of response. Percentage of up-regulated genes (Orange headers) or down-regulated genes (Blue headers) are calculated for each module, the highest percentages of response are used for visualization (Green headers).

**Figure 2.**
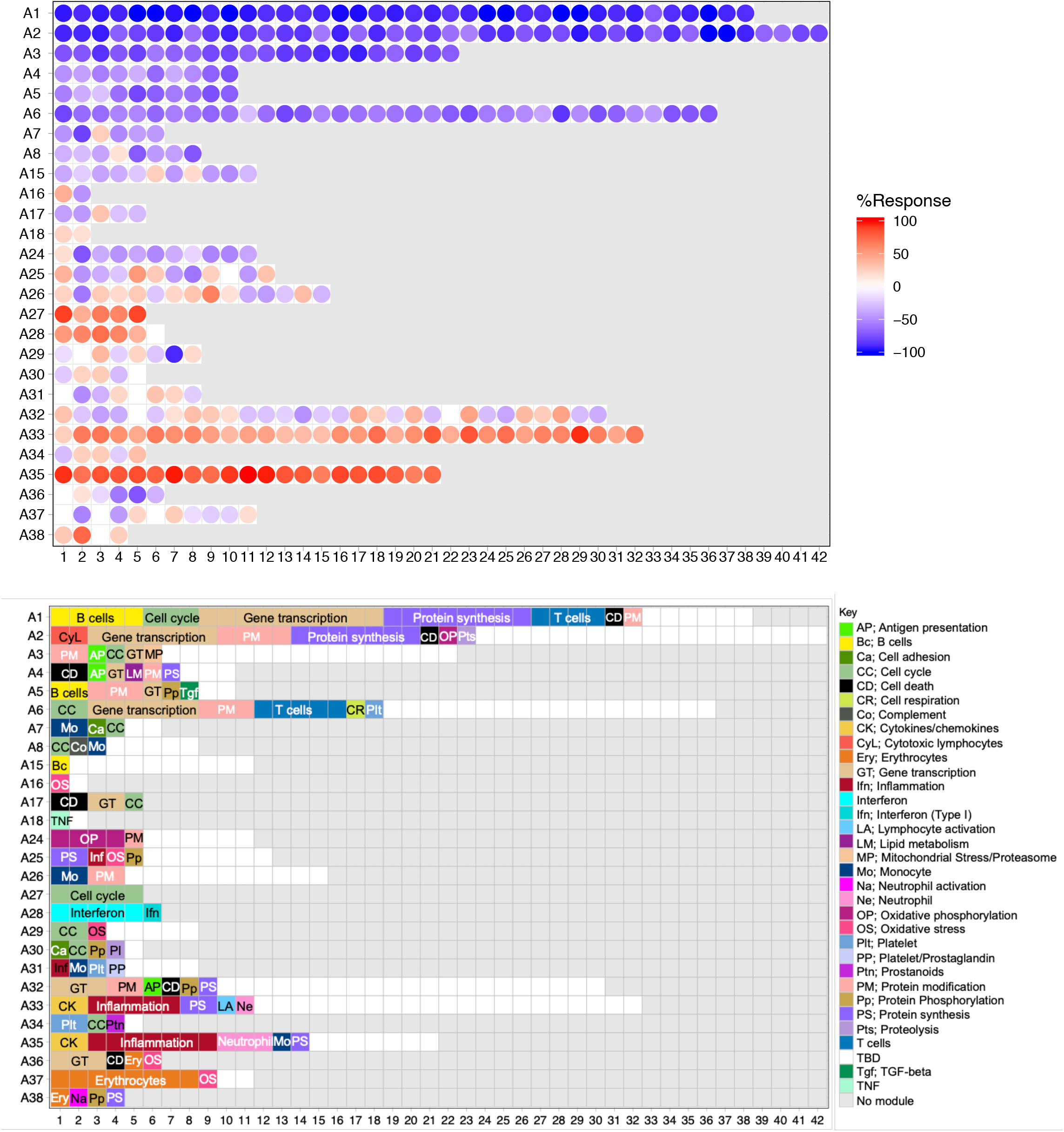
Module fingerprint grid plot. Each module is assigned a fixed position on a grid, with each row corresponding to a “module aggregate” constituted of modules following similar patterns of changes in abundance across 16 reference datasets corresponding to distinct immune states. The number of constitutive modules for each aggregate varies between 2 (A16 & A18) and 42 (A2). Red spots indicate modules for which constitutive transcripts are predominantly increased in septic patients in comparison to uninfected controls. This is expressed as a percent value representing the proportion of genes for the module in question which abundance is significantly increased. Conversely, blue spots indicate modules for which constitutive transcripts are predominantly decreased. The color key at the bottom indicate functions which have been attributed to some of the modules shown on the grid.

**Figure 3.**
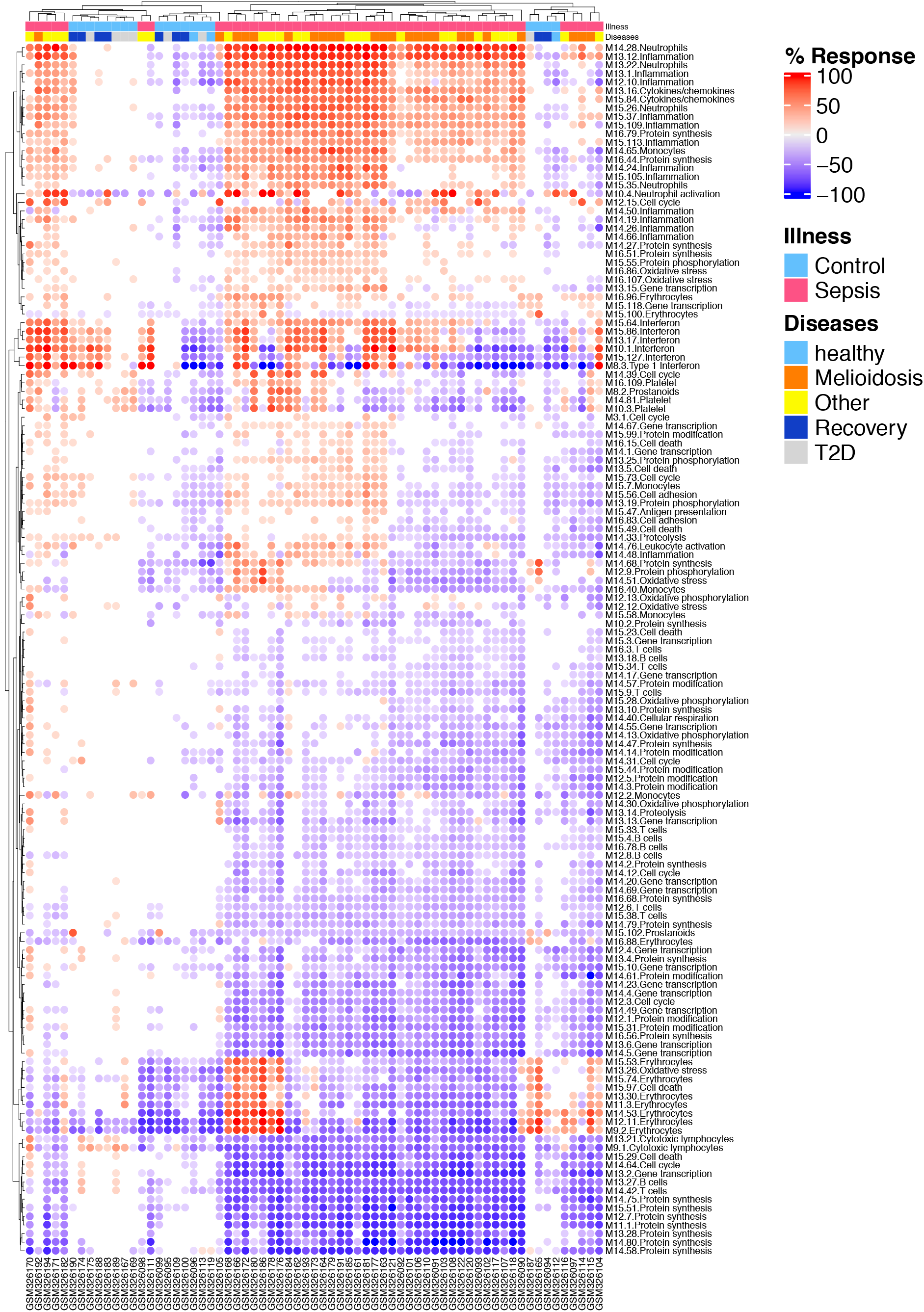
Fingerprint heatmap displaying patterns of annotated modules across individual study subjects. Fingerprint heatmap displaying patterns of differential transcript abundance in annotated modules across individual study subjects. The heatmap displays abundance patterns of 141 annotated modules for which at least 20% of constitutive genes were differentially expressed in at least one subject. Both samples (columns) and modules (rows) were arranged via hierarchical clustering. Proportion of differentially expressed transcripts is indicated by a color gradient ranging from blue (100% of transcript decreased) to red (100% of transcripts increased). Illness/Diseases information is indicated at the top of the heatmap using color codes.

### Step1: Annotation of the gene expression data matrix with module membership information

This step consists in assigning module membership information to each of the gene constituting the gene expression matrix.

The following R scripts have been developed for this step:

- “*1.2.1.Module_Matrix_preparation_datamatrix_byGEOquery.R*”. This script is compatible with datasets loaded using the “GEOquery” package (“*1.1Load_GEOdataset.R*”)
- “*1.2.2.Module_Matrix_preparation_datamatrix_byGEOmanually.R*”. This script is compatible with datasets which are downloaded manually from GEO (each table needs to be loaded to R).
- “*1.2.3.Module_Matrix_preparation_datamatrix_byCustomdata.R*”. This script is to be employed while using in-house generated datasets of any types (microarray, RNAseq etc…).

For our use case the input files for these scripts are:

- GSE13015 expression matrix (**Supplementary table 1**; which can also be obtained by running the script described in the previous section);
- Probe annotation table (**Supplementary table 2**)
- Sample information table (**Supplementary table 3**)
- Genes constituting the third version of blood transcriptome module repertoire (**Supplementary table 4** or “DC_ModuleGen3_2019.Rdata”).

The output file (**Figure 1**: step 1 = Annotation of the expression matrix) for this script is provided on **Supplementary table 5**. The output tables are split into 2 different files consisting of the data matrix table (“dat.mod.Gen3”) and the gene module annotation table (“dat.mod.func.Gen3”), which will be used in future steps. When running the script for datasets other than the one provided for the use case, changes should be made to the script for the “data matrix” loading, “probe annotation” and “sample information” steps.

### Steps 2&3: Differential expression and calculation of percentage response

These steps consist in performing, first group comparisons (Group1 vs control, Group 2 vs control) or individual sample comparisons (individual samples comparison to a control group or kinetic samples comparison to baseline control) and, secondly, computing for each module the proportion of constitutive transcripts for which abundance differs significantly between the study groups or individual single samples. An important point to consider at this step is the approach chosen for the selection of differentially expressed transcripts since it can be dependent on experimental design or preference in the choice of statistical cutoffs and multiple testing correction. For example, for this illustrative use case, we compared sepsis and control groups using *un-paired t-test*. The data were log2 transformed (assume log normalization) for differential expression analysis.

The following R scripts have been developed for this step:

- “*1.3.1.Statistic_for_Group_comparison.R*” for group comparison
- “*1.4.1.Statistic_for_Individual_analysis.R*” for single sample comparisons

Input files for this script are:

- Gene module matrix: “dat.mod.Gen3”
- Sample information: = “sample_info”
- Gene module annotation table: “dat.mod.func.Gen3”

Expected output files are:

- The percentage of response for group comparison: “res.mod.group”
- The percentage of response for individual comparison: “sum.mod.sin”

### Step 4: Grid and heatmap fingerprint visualization

The fourth step consists in visualizing changes in transcript abundance at the module-level. What was referred to earlier as the “module response” is expressed as the percentage the constitutive transcripts of a given module showing significant differential expression between study groups (e.g. case vs control, treated vs non-treated, pre-/post-treatment). Activity of each module is visualized as a red or blue spot which depicts percentage of transcripts which expression is increased or decreased, respectively. These spots can be arranged in a grid format, where modules occupy a fixed position on the grid. In the default grid configuration shown in **Figure 2**, modules on a given row belonged to one of 38 “module aggregates”. The modules aggregates were formed based on similarities in the patterns of transcript abundance changes across sixteen reference datasets used for construction of the modular repertoire. Thus, when the grid is read vertically across the rows it gives an indication of the patterns of change in a given set of patients at the least granular “module aggregate”-level. When the grid is read horizontally within each row and across the columns it gives an indication of the changes taking place at the more granular module-level. Functional interpretations are indicated by a color code that is overlaid on the grid plot (**Figure 2)**. On this grid module aggregates (rows) comprise between two and 43 modules (columns). Additional “aggregates” identified via clustering as outliers and comprising only one module are not shown on this grid, which, as a result is formed of only 27 rows (from a total of 38 aggregates). Because the positions of the modules on the grid are fixed, different fingerprints generated for independent groups of patients can be compared directly.

Changes in transcript abundance measured at the module-level can also be represented in a heatmap format with modules as rows and samples as columns, which can be rearranged based on similarities in module activity. The heatmap visualization can include all 382 modules or a pre-specified subset (e.g. based on functional annotation or aggregate information).

The following R scripts have been developed for module “fingerprint” visualization:

- “*1.3.2.Visualization_Grid_plot.R*” for Grid plot fingerprint visualization
- “*1.4.2.Visualization_Fingerprint_heatmap.R*” for fingerprint heatmap visualization

Input files for this script are:

- Group comparison matrix: “res.mod.group” from the previous step
- Individual comparison matrix: “sum.mod.sin” from the previous step

Expected output are images in JPEG and PDF formats shown on **Figure 2** (Group comparison) and **Figure 3** (Individual comparison).

## Discussion

Modular repertoires developed and by us and by others have been widely used for the analysis of blood transcriptome data. We released a first blood modular repertoire in 2008, and a second in 2013 (9,15,18). Notably, another blood modular repertoire has been developed by Li et al (19). The R script that is being shared here was developed to support analyses using the third iteration of transcriptional modules that we most recently developed. Compared to our earlier iterations, this new modular repertoire relied on an expended range collection of immune states, encompassing 16 datasets, corresponding to 16 distinct pathological or physiological immune states (16). The intent is for this transcriptional module repertoires to serve as a pre-established and stable framework for blood transcriptome data analysis. As such, rather than forming transcriptional modules for each new dataset that is being generated and analyzes, this same pre-established set of modules to be re-used for each new analysis. One of the advantages is the extended breadth of repertoire obtained when basing module construction on co-expression observed in a wider collection of immunological states, as opposed to doing so in a more unidimensional dataset (e.g. focusing on only one pathology). But maybe more importantly, because the repertoire is to be re-used over the years it is possible to dedicate much more time towards its functional characterization. This is illustrated by the depth of annotations provided for this set of modules, and which are accessible via the interactive presentations listed in **Table 1**. These annotations and results of functional profiling analyses are augmented by expression profiles of genes forming each module generated for different reference datasets.

Analytical tools and interactive web applications supporting module-level analyses and representations have previously been developed and disseminated, but only partially address user needs. Notably, a module enrichment web application and R package has been made available by January Weiner from the Max Planck institute in Berlin [insert as reference: (9,15,18) https://cran.r-project.org/web/packages/tmod/index.html].

The web-based tool supports blood transcriptome modules (both originating from the work of Li et al and ours) as well as other gene set collections compiled by MSigDB, the Molecular Signature Database hosted at the broad institute (20); http://software.broadinstitute.org/gsea/msigdb/genesets.jsp. Another high point of tmod is the implementation of statistical tests which are well suited for gene set-based analyses, including U-tests, hypergeometric tests as well customized approach developed by the same team [CERNO (21)]. However, the tool, logically, does not yet support the third iteration of blood transcriptome modules that we are just releasing (16). And it does not permit to generate fingerprints grids or perform module analysis at the individual sample level. We have in the past also developed web applications, which were employed not as analysis tools but as companions to earlier publications. These applications were designed as interactive supplements, giving readers the ability to adjust parameters and customize plots (e.g. change group comparisons or methods employed for multiple testing correction, “drill in” to gene-level data) (15,18). A wiki was also setup to consolidate information gathered as part of functional annotation / interpretation efforts. However, hosting of these resources was eventually discontinued, as dedicating funds to permit open-ended access post-publication is generally not a viable model for research organizations (this phenomenon is well documented in biomedical research and has been referred to as “url decay (22,23)). Since we have re-started efforts towards development of such resources. With recently applications deployed using R/Shiny that support interactive generation of fingerprint plots. These applications have been made available as ad hoc supplement to different publications (reference dataset for module construction: https://drinchai.shinyapps.io/dc_gen3_module_analysis/#; meta-analysis of respiratory syncytial virus blood transcriptome data: https://drinchai.shinyapps.io/RSV_Meta_Module_analysis/; development of targeted Covid-19 blood transcript panels: https://drinchai.shinyapps.io/COVID_19_project/). We will next undertake efforts to develop R packages supporting third party deployment of such applications. Notably, making here scripts available via GitHub and deploying apps using R-shiny should serve to future-proof these resources by guaranteeing availability on the longer term.

In the meantime, the R scripts which are provided, as part of this publication should help address existing gap “analytic gaps” when it comes to module repertoire analyses. They will permit the implementation of the data analysis and visualization workflow illustrated in recent use cases. This include notably a meta-analysis conducted starting from a collection of public blood transcriptome datasets comprised of profiles of subjects with respiratory syncytial virus (RSV) infection and uninfected controls (24). Blood repertoire fingerprints generated for each study at the group level were subsequently benchmarked. This permitted the identification of conserved modular signatures of RSV infection across studies. Conversely, fingerprints generated at the individual level permitted the identification of distinct molecular phenotypes shared among patients from different studies that were associated with clinical severity. This proof of principle illustrates potential utility for the workflow that is being supported by these R scripts while conducting integrative analyses.

However, important limitations remain. First it may be worth pointing out that the present resource is dedicated to the analysis and interpretation of human blood transcriptome data. It would thus not be suitable for analysis of data derived from other tissues or cell types. The only exception might be the investigation of immune infiltrates, for instance in tumor tissues, although performance is not guaranteed and one would need to proceed with caution with the interpretation of the data. Of note, we have participated in the development of module repertoires for mouse tissues, including blood (25). We also previously developed module repertoires based on co-expression observed in in vitro responses to immune stimuli (26,27). The script provided here could be employed for performing group and individual level module repertoire analyses and visualize results as heatmaps. But grid visualization would require a custom layout. However, it should be possible for users proficient in R programming language to adapt the script to handle visualization of other repertoires.

Regarding the analysis workflow itself, users may decide to adapt the script to fit their preferences for determining differential expression. It would be the case for both use of statistics and testing corrections. Also the script provided also assumes that pre-processing has been applied to normalize the data and that they have been Log2 transformed.

Other limitations are inherent to the module repertoire framework itself. For instance, our latest iteration still relies on microarray while RNAseq could have probably captured additional genes (likely resulting in larger sets of genes – whether modules could have been entirely missed is a matter of conjecture). The range of immune states covered by the sixteen input datasets is surely broader than that used in previous iterations but could yet be more comprehensive. These points are discussed in more details in the paper describing the construction of this framework (16). Importantly, the framework and scripts provided here have been found to be suitable for the analysis of various microarray platforms but also RNAseq data [e.g. (28)].

Another notable point is that the annotations shown on the grid are likely to evolve over time as more interpretation efforts continue. Also, it is recommended to check material deposited on GitHub for updates on a regular basis. The development of a web application supporting module repertoire analysis and visualization would be the next logical step for us and help address such shortcomings. For the time being we hope that the scripts that are being made available here will help at least partially address an unmet need regarding custom module repertoire analysis and visualization of blood transcriptome data.

## Availability of data and material

Scripts and data necessary to replicate analyses presented herein are shared on **GitHub;** https://github.com/Drinchai/DC_Gen3_Module_analysis.

A permanent DOI for this GitHub release was generated via Zenodo: https://zenodo.org/record/3944480

## Authors’ contributions

DR and DC conceptualization. DR, JR, WH, DB, DC: data curation and validation. DR, JR and DC: visualization. DR, JR, DC: analysis and interpretation. DB, DC: funding acquisition. DR and DC: methodology development. DR and DC: writing of the first draft. DR, JR, WH, DB, DC writing–review and editing The contributor’s roles listed above follow the Contributor Roles Taxonomy (CRediT) managed by The Consortia Advancing Standards in Research Administration Information (CASRAI) (https://casrai.org/credit/).

## References

1. Karsten SL, Kudo LC, Bragin AJ. Use of peripheral blood transcriptome biomarkers for epilepsy prediction. Neurosci Lett. 2011 Jun 27;497(3):213–7.

2. Freedman JE, Vitseva O, Tanriverdi K. The role of the blood transcriptome in innate inflammation and stroke. Ann N Y Acad Sci. 2010 Oct;1207:41–5.

3. Chaussabel D. Assessment of immune status using blood transcriptomics and potential implications for global health. Semin Immunol. 2015 Feb;27(1):58–66.

4. Pascual V, Chaussabel D, Banchereau J. A genomic approach to human autoimmune diseases. Annu Rev Immunol. 2010;28:535–71.

5. Mejias A, Suarez NM, Ramilo O. Detecting specific infections in children through host responses: a paradigm shift. Curr Opin Infect Dis. 2014 Jun;27(3):228–35.

6. Nakaya HI, Pulendran B. Vaccinology in the era of high-throughput biology. Philos Trans R Soc Lond B Biol Sci. 2015 Jun 19;370(1671).

7. Pascual V, Allantaz F, Patel P, Palucka AK, Chaussabel D, Banchereau J. How the study of children with rheumatic diseases identified interferon-alpha and interleukin-1 as novel therapeutic targets. Immunol Rev. 2008 Jun;223:39–59.

8. van Dam S, Võsa U, van der Graaf A, Franke L, de Magalhães JP. Gene co-expression analysis for functional classification and gene–disease predictions. Brief Bioinform. 2017 Jan 10;19(4):575–92.

9. Chaussabel D, Quinn C, Shen J, Patel P, Glaser C, Baldwin N, et al. A modular analysis framework for blood genomics studies: application to systemic lupus erythematosus. Immunity. 2008 Jul 18;29(1):150–64.

10. Ardura MI, Banchereau R, Mejias A, Di Pucchio T, Glaser C, Allantaz F, et al. Enhanced monocyte response and decreased central memory T cells in children with invasive Staphylococcus aureus infections. PloS One. 2009;4(5):e5446.

11. Banchereau R, Jordan-Villegas A, Ardura M, Mejias A, Baldwin N, Xu H, et al. Host immune transcriptional profiles reflect the variability in clinical disease manifestations in patients with Staphylococcus aureus infections. PloS One. 2012;7(4):e34390.

12. Berry MPR, Graham CM, McNab FW, Xu Z, Bloch SAA, Oni T, et al. An interferon-inducible neutrophil-driven blood transcriptional signature in human tuberculosis. Nature. 2010 Aug 19;466(7309):973–7.

13. Tattermusch S, Skinner JA, Chaussabel D, Banchereau J, Berry MP, McNab FW, et al. Systems biology approaches reveal a specific interferon-inducible signature in HTLV-1 associated myelopathy. PLoS Pathog. 2012 Jan;8(1):e1002480.

14. Quartier P, Allantaz F, Cimaz R, Pillet P, Messiaen C, Bardin C, et al. A multicentre, randomised, double-blind, placebo-controlled trial with the interleukin-1 receptor antagonist anakinra in patients with systemic-onset juvenile idiopathic arthritis (ANAJIS trial). Ann Rheum Dis. 2011 May;70(5):747–54.

15. Obermoser G, Presnell S, Domico K, Xu H, Wang Y, Anguiano E, et al. Systems scale interactive exploration reveals quantitative and qualitative differences in response to influenza and pneumococcal vaccines. Immunity. 2013 Apr 18;38(4):831–44.

16. Altman MC, Rinchai D, Baldwin N, Toufiq M, Whalen E, Garand M, et al. Development and Characterization of a Fixed Repertoire of Blood Transcriptome Modules Based on Co-expression Patterns Across Immunological States. bioRxiv. 2020 Apr 28;525709.

17. Pankla R, Buddhisa S, Berry M, Blankenship DM, Bancroft GJ, Banchereau J, et al. Genomic transcriptional profiling identifies a candidate blood biomarker signature for the diagnosis of septicemic melioidosis. Genome Biol. 2009;10(11):R127.

18. Chaussabel D, Baldwin N. Democratizing systems immunology with modular transcriptional repertoire analyses. Nat Rev Immunol. 2014;14(4):271–80.

19. Li S, Rouphael N, Duraisingham S, Romero-Steiner S, Presnell S, Davis C, et al. Molecular signatures of antibody responses derived from a systems biology study of five human vaccines. Nat Immunol. 2014 Feb;15(2):195–204.

20. Liberzon A, Birger C, Thorvaldsdóttir H, Ghandi M, Mesirov JP, Tamayo P. The Molecular Signatures Database (MSigDB) hallmark gene set collection. Cell Syst. 2015 Dec 23;1(6):417–25.

21. Zyla J, Marczyk M, Domaszewska T, Kaufmann SHE, Polanska J, Weiner J. Gene set enrichment for reproducible science: comparison of CERNO and eight other algorithms. Bioinforma Oxf Engl. 2019 15;35(24):5146–54.

22. Wren JD. URL decay in MEDLINE--a 4-year follow-up study. Bioinforma Oxf Engl. 2008 Jun 1;24(11):1381–5.

23. Wren JD, Georgescu C, Giles CB, Hennessey J. Use it or lose it: citations predict the continued online availability of published bioinformatics resources. Nucleic Acids Res. 2017 20;45(7):3627–33.

24. Rinchai D, Konza O, Hassler S, Martina F, Mejias A, Ramilo O, et al. Characterizing blood modular transcriptional repertoire perturbations in patients with RSV infection: a hands-on workshop using public datasets as a source of training material. bioRxiv. 2019 Jan 22;527812.

25. Singhania A, Graham CM, Gabryšová L, Moreira-Teixeira L, Stavropoulos E, Pitt JM, et al. Transcriptional profiling unveils type I and II interferon networks in blood and tissues across diseases. Nat Commun. 2019 28;10(1):2887.

26. Alsina L, Israelsson E, Altman MC, Dang KK, Ghandil P, Israel L, et al. A narrow repertoire of transcriptional modules responsive to pyogenic bacteria is impaired in patients carrying loss-of-function mutations in MYD88 or IRAK4. Nat Immunol. 2014 Dec;15(12):1134–42.

27. Banchereau R, Baldwin N, Cepika A-M, Athale S, Xue Y, Yu CI, et al. Transcriptional specialization of human dendritic cell subsets in response to microbial vaccines. Nat Commun. 2014 Oct 22;5:5283.

28. Rawat A, Rinchai D, Toufiq M, Marr A, Kino T, Garand M, et al. A neutrophil-driven inflammatory signature characterizes the blood cell transcriptome fingerprints of Psoriasis and Kawasaki Disease. bioRxiv. 2020 Feb 25;2020.02.24.962621.

